# Environmental DNA for the enumeration and management of Pacific salmon

**DOI:** 10.1101/394445

**Authors:** Taal Levi, Jennifer M. Allen, Donovan Bell, John Joyce, Joshua R. Russell, David A. Tallmon, Scott C. Vulstek, Chunyan Yang, Douglas W. Yu

## Abstract

Pacific salmon are a keystone resource in Alaska, generating annual revenues of well over ∼US$500 million/yr. Due to their anadromous life history, adult spawners distribute amongst thousands of streams, posing a huge management challenge. Currently, spawners are enumerated at just a few streams because of reliance on human counters and, rarely, sonar. The ability to detect organisms by shed tissue (environmental DNA, eDNA) promises a more efficient counting method. However, although eDNA correlates generally with local fish abundances, we do not know if eDNA can accurately enumerate salmon. Here we show that daily, and near-daily, flow-corrected eDNA rate closely tracks daily numbers of returning sockeye and coho spawners and outmigrating sockeye smolts. eDNA thus promises accurate and efficient enumeration, but to deliver the most robust numbers will need higher-resolution stream-flow data, at-least-daily sampling, and a focus on species with simple life histories, since shedding rate varies amongst jacks, juveniles, and adults.

## Introduction

Pacific salmon (*Oncorhynchus* spp.) support a $449 million/yr commercial fishery, play a significant role in the $470 million/yr sport fishery (National Marine Fisheries Service 2017) in Alaska alone, and remain a key cultural and subsistence resource for humans. Salmon are also a major source of marine nutrient and energy subsidies to terrestrial and aquatic food webs, in large part by being important seasonal prey resources for bears, eagles, and other culturally, biologically, and economically important consumers (Gende *et al.* 2002; Gende *et al.* 2004; Schindler *et al.* 2003; Shakeri *et al.* 2018; Wheat *et al.* 2017). Due to their anadromous life history, salmon fisheries are often managed by setting escapement goals, where escapement refers to the number of fish that escape the mostly ocean-based fishery and are thus available for spawning in fresh water. For example, from April to October each year, the Alaska Department of Fish and Game (ADFG) continuously estimates salmon breeding population sizes in some Alaskan streams and issues temporary fishery closure notices to ensure that these escapements exceed minimum target sizes per species.

Of course, it is very costly to count fish. A typical salmon weir consists of a series of closely spaced bars across an entire stream to prevent the passage of salmon, except through a single, narrow gate over which a human observer tallies and identifies to species salmon as they file through (alternatively, Didson sonar can be used to count and size salmon individuals as they pass with species identity inferred from body size and run timing). The annual operating cost of a weir is approximately $80,000, not including installation or major maintenance (Fox 2018), and even this setup might be prone to undercounting (Eggers *et al.* 2009).

More than 6,000 streams are used by various combinations of the five species of Pacific salmon in Southeast Alaska alone, and more than 1000 of those streams have been documented as hosting spawning populations (Johnson & Blossom 2018; Fig. S1). Not surprisingly, almost all these salmon runs are left unmonitored or are monitored only every few years with crude indices such as visual transects conducted on foot or from the air. Detailed sampling effort varies depending upon budgets, but only a few streams are enumerated and are given escapement targets in any given year. For example, coho salmon (*O. kisutch*) are managed in Southeast Alaska by monitoring escapements and commercial fishery take from only four to nine full indicator stock streams (Shaul *et al.* 2005). Full indicator stock streams are those in which juveniles (usually outmigrating smolts) are tagged with coded wire tags and marked with an adipose fin clip. The proportion of marked fish sampled upon return, along with fishery and escapement sampling, are used to estimate smolt production, fishery interception rate, and escapement. Additional coho streams near urban centers are surveyed by air or on foot, and in some cases escapement goals are established, but there is no guarantee that these intermittent surveys overlap with the peak abundances of runs. Similarly, sockeye salmon (*O. nerka*) escapements are at least partially enumerated at only fourteen streams in Southeast Alaska (Munro & Volk 2016). Nearly all pink (*O. gorbuscha*) and chum (*O. keta*) salmon runs are left un-enumerated by weirs or sonar, despite these species making up the majority of salmon biomass, harvest, and economic value in this region. Instead, several larger chum and pink streams are surveyed by air or on foot several times each year (Munro & Volk 2016), but even this is complicated by the difficulty of distinguishing pink and chum because their migration timing and habitat use often overlap. Finally, enumeration is naturally focused on the largest, most economically valuable streams, leaving large numbers of subdominant runs for most salmon species unmonitored most years.

Fry and smolt production resulting from spawning salmon is monitored with even less effort, which limits inference of future expected recruitment and harvest. Poor understanding of fry and smolt production also limits inference regarding the degree to which salmon productivity is limited by spawning habitat for adults or by rearing habitat for juveniles, and whether changes in marine or freshwater productivity are responsible for changes in salmon recruitment and abundance. Such information is critical for informed management and for judging the potential efficacy of stock enhancement programs.

More generally, the under-monitoring of Pacific salmon stocks hinders the construction of reliable spawner-recruit models, which are used to determine escapement goals for maximum sustainable yield. The lack of such models increases uncertainty about whether, and where, there are sufficient spawners to maximize recruitment and increases the risk of long-term decline or loss, especially of the small, subdominant components of salmon runs. These smaller salmon runs increase the resilience of salmon stocks through portfolio effects (Schindler *et al.* 2010), can restock a dominant component that has suffered a negative shock, and provide key resources for wildlife by extending the spatial range and phenology of salmon availability to terrestrial and aquatic food webs (Gende *et al.* 2002; Levi *et al.* 2015; Schindler *et al.* 2013). As fisheries increasingly transition towards ecosystem-based fisheries management (Levi *et al.* 2012), identifying, monitoring, and maintaining such spatially and temporally distributed salmon resources becomes increasingly important for conservation and management.

The advent of environmental DNA (eDNA) methods that detect DNA shed by organisms (Bohmann *et al.* 2014; Goldberg *et al.* 2016) provides a promising tool for monitoring salmon escapements and juvenile production because it could increase management-relevant information at low cost. However, while the efficacy of using eDNA for species *detection* is now widely recognized (Goldberg *et al.* 2016; Rees *et al.* 2014) and while several studies have demonstrated that eDNA is generally correlated with fish abundance in mesocosm experiments, lakes, and streams (Doi *et al.* 2015; Handley *et al.* 2018; Lacoursière-Roussel *et al.* 2016; Takahara *et al.* 2013; Tillotson *et al.* 2018; Wilcox *et al.* 2016), we do not yet know whether eDNA contains sufficient information to robustly and accurately estimate fish abundance, particularly for anadromous fish as they enter and leave a watershed. By robust, we mean accuracy that is not greatly affected by variation among years, species, stream, and/or details of the sampling protocol.

Anadromous fish such as salmon provide a straightforward scenario for testing whether eDNA can be used to count fish, because potentially large numbers of salmon release their DNA as they pass a fixed sampling point, either as they swim upstream as returning adults or swim downstream as outmigrating juveniles. If eDNA degrades or settles quickly (as suggested by Jane *et al.* 2015; Jerde *et al.* 2016; Sassoubre *et al.* 2016; Shogren *et al.* 2016; Turner *et al.* 2015), then eDNA concentrations should primarily detect fish that are locally present in space and time. Thus, rather than simply accumulating as fish enter a watershed, eDNA concentrations might spike up and down as a pulse of fish swims past a sampling point, with the size of the spike correlated with fish number and/or biomass. Because the concentration of eDNA in streamwater results from both the amount of DNA shed by organisms and the flow of water, the product of eDNA concentration and streamflow (measured in units of water volume per time) can be used to calculate absolute quantities of eDNA per unit time. Such ‘flow-corrected eDNA rates’ measured at regular intervals (e.g. daily) could then be substituted for, or complement, gold-standard count data from weirs. For sockeye, coho, and chinook salmon, which produce juveniles that typically rear in freshwater prior to outmigrating to the ocean as smolts, whether this is plausible depends on the strength of eDNA signal produced by adults relative to what is produced by juveniles residing upstream. If, for instance, juveniles rear sufficiently far upstream, the signal of their eDNA should be weak or undetectable, eliminating a source of noise that would prevent the robust enumeration of adult salmon entering lower stream reaches with eDNA.

In the most comprehensive and relevant study to date, Tillotson *et al.* (2018) demonstrated that local counts of sockeye salmon in a spawning creek, particularly dead sockeye, indeed predict local eDNA concentrations. As Tillotson *et al.* (2018) put it, the next step is “reversing the model to predict abundance from eDNA.” We accomplish this by taking advantage of a daily census of sockeye and coho salmon carried out at the Auke Creek research weir in Juneau, Alaska to test whether eDNA concentrations and stream-flow measurements together produce quantitative and management-relevant indices of salmon escapement and juvenile outmigration. To explore the general ecology of eDNA, we also quantify the relative influences of salmon counts on the same day of water sampling, salmon that entered the watershed one day prior, and salmon that entered two days prior to an eDNA measurement, and we assess the eDNA signal produced by salmon of different life stages and body sizes. The purpose of these latter analyses is to test for two possible sources of error (long-distance transport of eDNA and differential shedding rates by body size and type) when using eDNA to enumerate salmon.

## Methods

### Weir operation

The Auke Creek research weir is located 19.2 km north of Juneau, Alaska, 400 m downstream from the outlet of Auke Lake above the high tide line at the mouth of Auke Creek (Fig. 1). The ∼1072.5 ha watershed includes five tributaries that feed into Auke Lake, which is 1.6 km long and 1.2 km wide, with a surface area of 67 ha. The weir is cooperatively operated by the National Marine Fisheries Service, in collaboration with the University of Alaska, and the Alaska Department of Fish and Game, with the objective of capturing all outmigrants and returning spawners at Auke Creek. All outmigrants (from upstream) are enumerated from the beginning of March to the middle of June and released below the weir, after which the weir is converted to capture returning adult salmonids (from downstream), which are counted and then released above the weir. During monitoring of adult salmonids, fish are classified by species and life stage. The Auke Creek dataset represents probably the highest-temporal-resolution and most accurate wild Pacific salmon census data in Alaska, if not the world. Life stages for coho salmon include typical adult male and female fish along with smaller early maturing and small-bodied ‘jack’ males, and a unique ‘nomadic’ juvenile life-history strategy in which coho fry rearing in the estuary and ocean return upstream (Koski 2009). Coho ‘nomads’ are similar to ocean-type chinook and sockeye salmon that outmigrate as fry rather than rearing in freshwater, with the exception that coho ‘nomads’ rear in the estuary within their salt tolerance and return to freshwater in the fall where they overwinter as juveniles before outmigrating to the ocean as smolts the following year (see Koski 2009 for details). Sockeye salmon can also produce jacks, but infrequently. Complete methods for weir operation can be found in Vulstek *et al.* (2018) (Weir photos in Supplemental information S2). River height is recorded daily and converted to streamflow (cubic feet per second) using an established rating curve (Bell *et al.* 2017).

**Figure 1.**
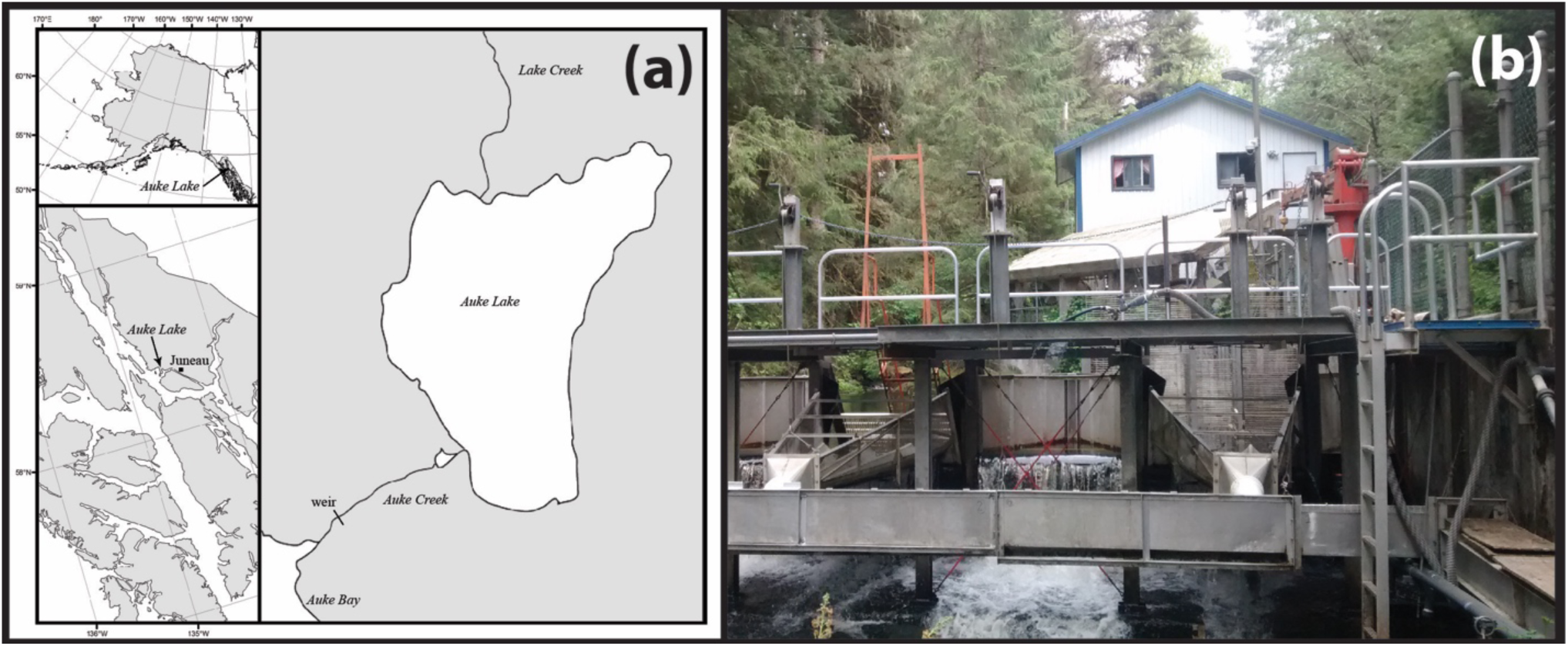
The Auke Creek research weir is (A) located in Juneau, Alaska at the outflow of Auke Lake. (B) The weir is a permanent structure used to sort and enumerate outmigrating juvenile salmon and returning adult salmon.

### Environmental DNA quantitation

We collected water samples from just upstream of the weir (location photograph in Supplemental Information S2) for three years, from 2014-2016, after each day’s salmon enumeration. In a 2014 pilot study, we collected three 1L water samples weekly from 28 May to 11 December. Based on promising results, and to reduce costs, in 2015, we sampled weekly when few fish were entering the river and then increased sampling frequency up to daily during periods in which many salmon were entering the river. Because salmon eDNA disappeared entirely after October in 2014, we sampled from 12 May to 3 November in 2015. Based on further promising results from 2015, we increased sampling frequency to daily in 2016 from 10 May to 20 October. Because previous technical replicates had yielded consistent results, and because of the high frequency of water collection, we collected only two 1L water samples daily in 2015 and 2016. All water samples were collected using 1L disposable sterile Whirlpak bags and filtered through a 0.45 micron cellulose nitrate filter. Filters were then folded and stored in 100% ethanol at 4C until laboratory processing.

We maintained strict protocol to prevent contamination of filters and reagents. We performed DNA extraction and PCR setup inside of separate HEPA-filtered and UV-irradiated PCR cabinets (Air Science LLC, Fort Meyers, FL) within a separate lab where PCR product is prohibited. Filters were first removed from ethanol and air-dried overnight in sterile, disposable weigh boats. A modified protocol for the Qiagen DNeasy Blood and Tissue kit was used to isolate DNA. This included the addition of 1.0 mm zirconia/silica beads to the initial lysis buffer and then a 15 minute vortex step to loosen the DNA from the filters. Incubation in lysis buffer was increased to 48 hours. After incubation, 300 ul of the lysed product was transferred to a new 1.7 ml microcentrifuge tube. Thereafter, we followed the manufacturer’s protocol. DNA was eluted in a total volume of 100 ul.

Using species-specific primers and TaqMan minor groove binder (MGB) probes (ThermoFisher Scientific, Waltham, MA), developed by Rasmussen Hellberg *et al.* (2010) (Table 1), we targeted a fragment of the cytochrome c oxidase subunit 1 (COI) gene. For each species, each sample was run in triplicate PCRs. Each 20 ul qPCR contained 6 ul of DNA template, 10 ul Environmental Master Mix 2.0 (ThermoFisher Scientific, Waltham, MA), 0.2 uM of both forward and reverse primers, 0.2 um of the TaqMan MGB probe, and sterile water. Additionally, each plate contained a four-point standard curve using DNA obtained from salmon tissue from each species. Extracted tissue was quantified using a Qubit Fluorometer (ThermoFisher Scientific, Waltham, MA) and diluted 10-fold from 10-1 to 10-4 ng/ul. PCR cycling conditions involved an initial denaturation step of 10 min at 95 °C to activate the HotStart Taq DNA polymerase, followed by 50 cycles of 95 °C for 15 s and 60 °C for 60 s. All reaction plates contained a negative control (water) as well as extraction blanks. PCR was performed on an ABI PRISM 7500 FAST Sequence Detection System (Applied Biosystems, Foster City, CA) and analyzed on 7500 Software v2.0.6 (Applied Biosystems, Foster City, CA). Cycle values were converted to target-DNA concentration using the standard curve derived from the tissue samples, and each day’s eDNA concentration was taken as the mean across the two extractions and the three qPCR replicates from that day for that species.

**Table 1.**
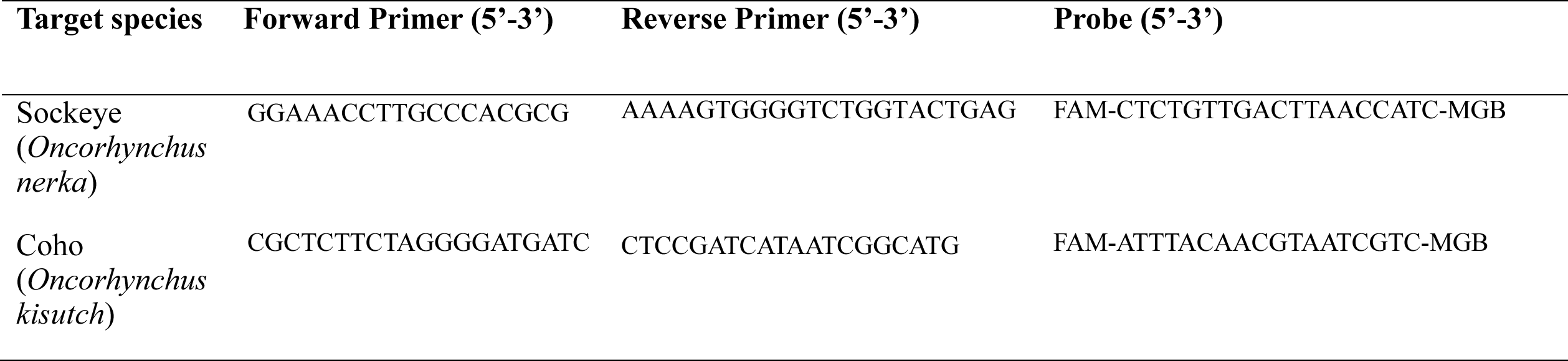
Species-specific primers and probes used in this study (Rasmussen Hellberg *et al.* 2010)

### Data analysis

To calculate the flow-corrected eDNA rate, we multiplied each day’s qPCR-estimated target-DNA concentration 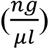 against that day’s streamflow 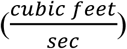. There is no need to harmonize units because the product is now an estimate of DNA biomass rate (ng/sec) multiplied by a dimensionless constant (volume/volume): 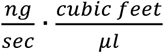, and the fitted model parameters incorporate the conversion factor. Streamflow was usually taken at 8 AM each day, near the time that eDNA was sampled. Note that this measure is only for one time point per day and might not be fully representative of streamflow over the whole day.

We predicted salmon counts from the natural log of flow-corrected eDNA rate using a quasipoisson regression with a log-link function in order to account for overdispersed count data. The quasipoisson model produces the same coefficients as standard Poisson generalized linear models for count data, but it is more inferentially conservative (i.e. lower Type I error rates due to wider confidence intervals). Log transformation of flow-corrected eDNA rate (1) allowed for the fit of zero salmon counts in the Poisson model, which would otherwise only be achievable if the eDNA rate approached negative infinity due to the log-link, and (2) fit a flexible power law (a linear model fit in log-log space). We fit separate models in 2015 and 2016 for returning adult sockeye salmon, returning total coho salmon, and outmigrating sockeye smolts. In our analysis, we included data for adult sockeye salmon from 18 June - 1 August, adult coho salmon from 15 August – 30 October, and outmigrating sockeye smolts from 15 April – 10 June. This time period captured the full runs of each species and life stage, but did not include a time period after the adult sockeye salmon run when DNA was transported downstream as salmon died in the lake. We used total coho, not just adult coho, because the coho run includes a varying mixture of nomadic juveniles, jacks, and adults, which are different sizes but with unknown relative contributions to DNA that we found to not scale predictably with biomass (see *Ecology of eDNA*). This is complicated in part by the fact that much larger-bodied adult salmon do not eat or defecate unlike juvenile nomads, potentially unlinking the rate of eDNA shedding to biomass or surface area. We detected a single high leverage outlier for coho salmon in 2016 in which a day with a large pulse of jacks retained a low concentration of eDNA. To avoid poor model predictions due to this outlier, we removed this data point from the results in the main text and include this outlier in the models in Supplemental Information S3.

To determine whether the relationship between flow-corrected eDNA and salmon counts was consistent between the two years, we combined the data from the two years and fit a model with an additional interaction term between year and flow-corrected eDNA. A significant interaction effect would indicate a different relationship between count and eDNA between years, which would indicate a lack of model transferability.

We collected daily water temperature data, but we observed a strong negative correlation of temperature and flow (*r* = −0.75 for Sockeye adult dataset, *r* = −0.53 for Sockeye smolt dataset, *r* = −0.10 for coho total dataset), and we were concerned about spurious correlations caused by the temporal trend in temperature. Nevertheless, we explored models with stream temperature and observed no consistent results in the magnitude, sign, or significance of the temperature effect across years or species, which suggested to us that our concern about spurious temperature effects were warranted and could lead to overfitting that exaggerated the precision of our predicted number of counts. That is, temperature effects were not transferable among years within the same salmon type or among salmon types.

### Ecology of eDNA

We also used the dataset to explore the ‘ecology of eDNA,’ using salmon counts from the same and previous days to predict that day’s flow-corrected eDNA rate. The purpose is to test for the possibility that long-distance, albeit attenuated, transport of eDNA from far-upstream salmon degrades the real-time quantitative accuracy of eDNA. We also test for the possibility that body size and/or life-history affects per-fish shedding rates.

To directly estimate the timescale over which eDNA was detected in Auke Creek, we used a series of three linear regression models to relate daily counts of sockeye salmon in 2016 (the year with daily sampling) to flow-corrected eDNA concentration. We first modeled flow-corrected eDNA as a function of salmon counts from the same day. We then used the residuals from that model in a second regression that instead included salmon counts from the previous day as a predictor. Finally, we used the residuals from the second model in a regression using salmon counts from two days prior as a dependent variable. We interpreted significant lag variables from salmon counts in the second or third models as evidence that salmon entering the river one or two days ago influence the measured flow-corrected eDNA concentration. In order to explore the eDNA production by coho salmon of different life stages, we additionally used multiple linear regression with counts of adults, jacks, and nomad juveniles in 2015 and 2016 as predictors of flow-corrected eDNA measured that same day.

## Results

Neither the concentration of eDNA nor flow-corrected eDNA rate increased monotonically as salmon accumulated in the Auke Creek watershed. Instead, flow-corrected eDNA rates reflected a highly local signal of salmon abundance in space and time, effectively tracking salmon that had passed near the water sampling site over the previous day (Figs. 2-4). This was true for both adult salmon and smolts.

**Figure 2.**
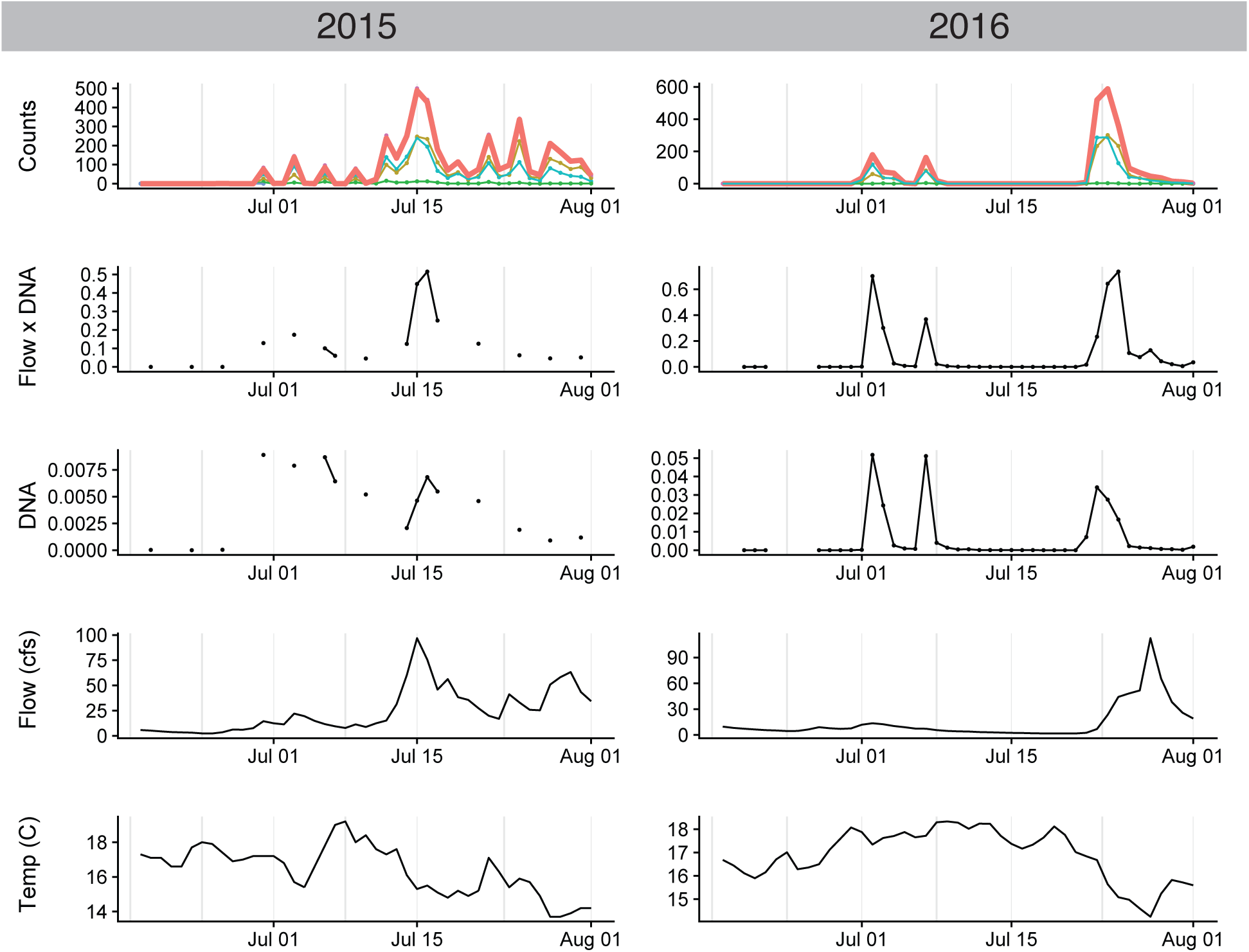
Timeline from June 18 to August 1 of adult sockeye salmon counts, flow-corrected eDNA concentration (ng/μl*cfs), uncorrected eDNA concentration (ng/μl), stream flow (cfs, cubic-feet/sec), and stream temperature (°C) in 2015 and 2016. Environmental DNA results from consecutive days are connected by lines. Male and female salmon are denoted by yellow-brown and blue lines respectively, and jacks are denoted by green lines. Total adult sockeye salmon counts are denoted by thick red lines.

**Figure 3.**
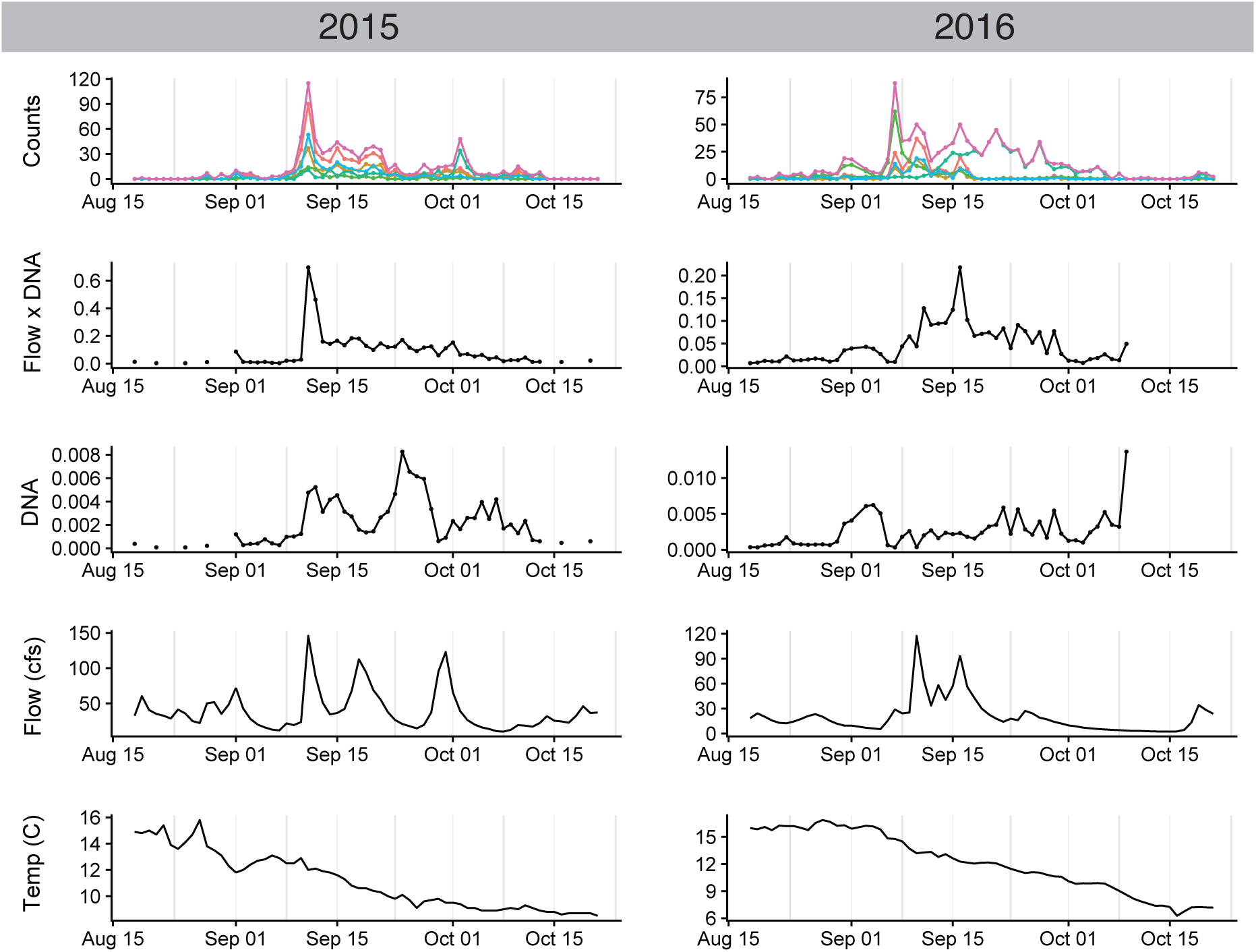
Timeline from August 15 to October 30 of coho salmon counts, flow-corrected eDNA concentration (ng/μl*cfs), uncorrected eDNA concentration (ng/μl), stream flow (cfs), and stream temperature (C) in 2015 and 2016. Environmental DNA results from consecutive days are connected by lines. Male and female coho salmon are denoted by yellow-brown and blue lines respectively, jacks are denoted by green lines, counts of a nomadic juvenile life history strategy in which young coho rear in the estuary and ocean and return upstream are denoted by teal lines, total adult (male + female) coho salmon counts are denoted by red lines. Total coho salmon counts including jacks and juveniles are denoted by pink lines. Note that the adult male and female coho salmon were the dominant component of the run in 2015 while the jack and juvenile life history strategy was a major component of the run in 2016. A pulse of 62 coho jacks was recorded on Sep 7, 2016, but no concomitant eDNA signal was recorded.

### Tracking of salmon phenology and abundances with eDNA

The natural logarithm of the product of stream flow (cubic feet per second, cfs) and eDNA concentration (ng/μl), which we refer to as flow-corrected eDNA rate, was highly predictive of the counts of returning adult sockeye and coho salmon, as well as of outmigrating sockeye salmon smolts in both 2015 and 2016 (Fig. 5; Adult sockeye 2015: β = 0.63±0.20, *p* = 0.008; Adult sockeye 2016: β = 0.79±0.11, *p* < 2e-8; Adult sockeye both years: β = 0.71±0.09, *p* < 2e-10; Total coho 2015: β = 0.70±0.10, *p* < 2e-8; Total coho 2016: β = 0.78±0.10, *p* < 3e-10; Total coho both years: β = 0.66±0.06, *p* < 2e-16; Sockeye smolts 2015: β = 1.64±0.37, *p* = 0.004; Sockeye smolts 2016: β = 1.42±0.35, *p* = 0.003); Sockeye smolts both years: β = 1.33±0.30, *p* = 0.0005).

**Figure 4.**
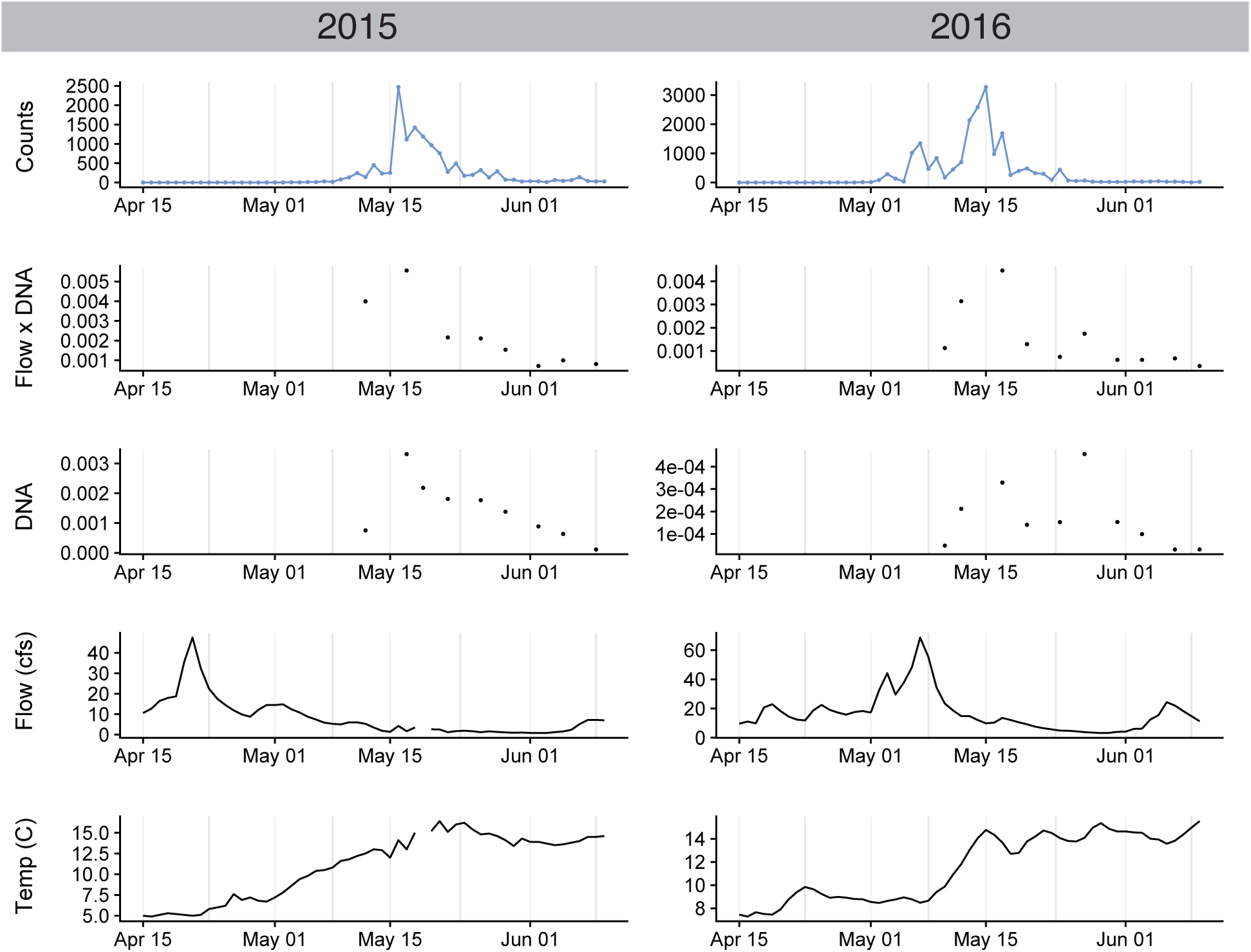
Timeline from April 15 to June 10 of outmigrating sockeye salmon smolt counts, flow-corrected eDNA concentration (ng/μl*cfs), uncorrected eDNA concentration (ng/μl), stream flow (cfs), and stream temperature (C) in 2015 and 2016. Environmental DNA results from consecutive days are connected by lines.

**Figure 5.**
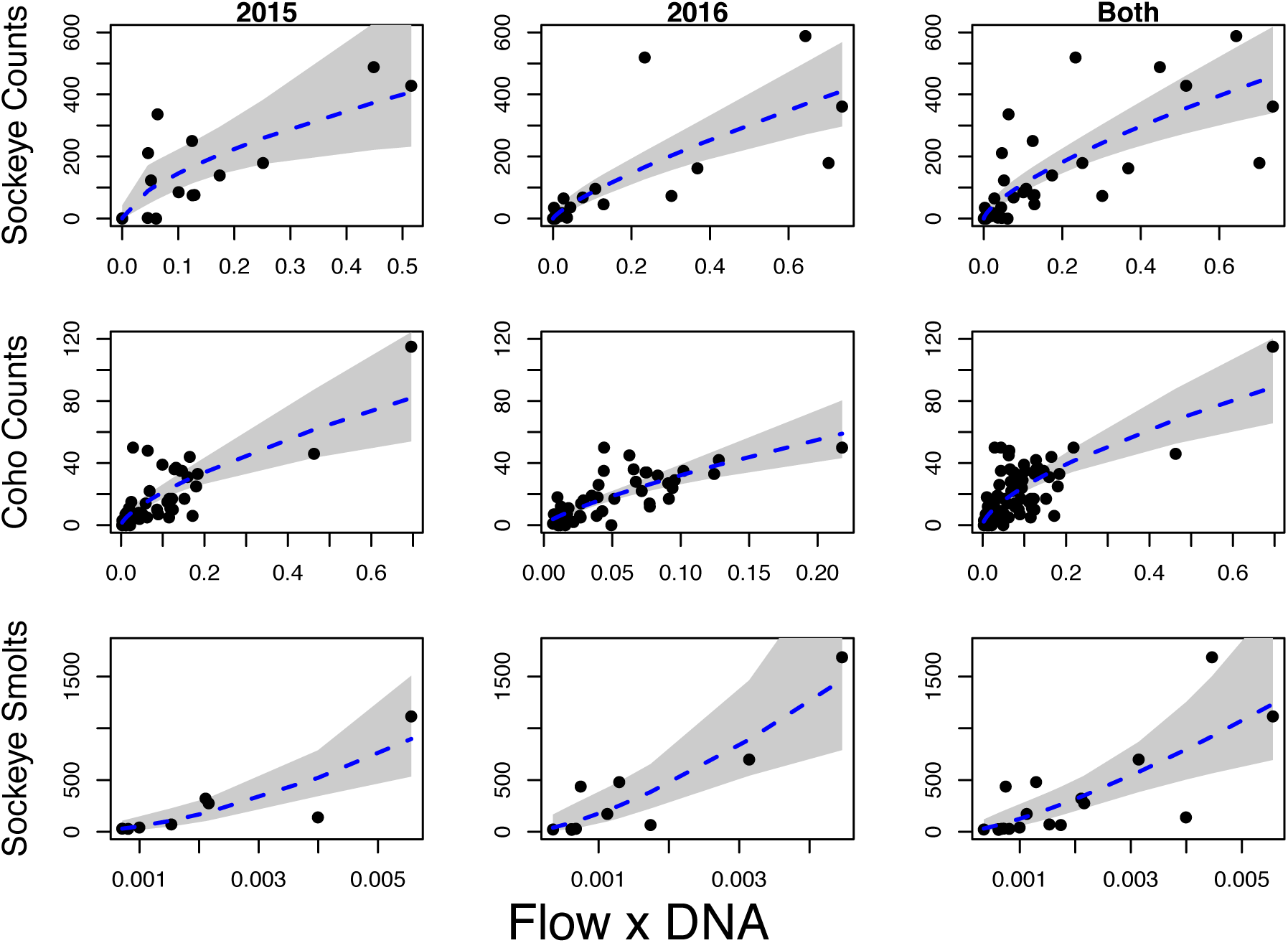
Results of quasipoisson regression models relating flow-corrected eDNA concentration to adult sockeye salmon counts (2015: p=0.008, 2016: p<2e-8, both years: p<1e-10), total coho salmon counts (2015: p<2e-8, 2016: p<3e-8, both years: p<2e-16), and counts of sockeye salmon smolts (2015: p=0.004, 2016: p=0.003, both years: p<0.005). Gray shading denotes the 95% confidence interval.

The combined models for 2015 and 2016 unambiguously failed to identify an interaction effect between year and flow-corrected eDNA rate for adult sockeye salmon (*p* = 0.43), total coho salmon (*p* = 0.59), and for sockeye salmon smolts (*p* = 0.71), indicating that eDNA had a consistent relationship with salmon counts across years.

In all models, the quasipoisson regression models using flow-corrected eDNA rate as a single predictor produced visually representative predictions of counts through time that captured the phenology, temporal dynamics, and relative abundance of each run (Fig. 6). Similarly, we tried models with water temperature as an additional predictor but saw no consistently significant effects, and we were concerned about spurious correlations caused by the temporal trend in temperature data and its strong anti-correlation with flow (*r* = −0.75 for Sockeye adult dataset, *r* = −0.53 for Sockeye smolt dataset, *r* = −0.10 for coho total dataset). This result is not surprising since visual inspection of the temperature timelines (Figs. 2-4) reveals no covariance with fish counts.

**Figure 6.**
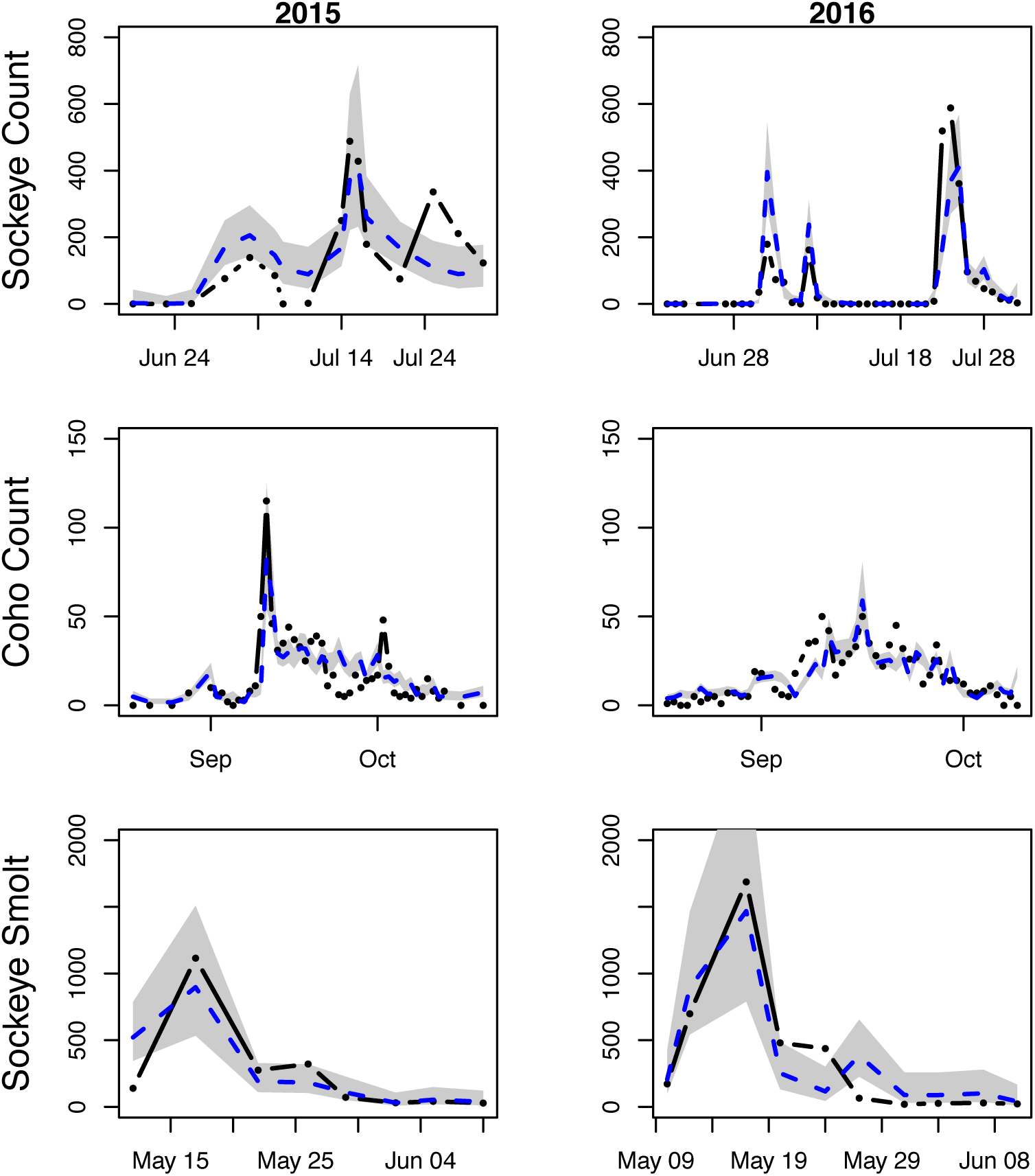
Counts of adult sockeye salmon, total coho salmon (including all life history strategies), and sockeye salmon smolts (black dots) and the predicted number of counts based on the flow-corrected eDNA concentration predictor in the quasipoisson regression model (blue dashed lines). Gray shading denotes the 95% confidence interval.

### Ecology of eDNA

As expected, sockeye salmon counts from the current day in 2016 significantly predicted flow-corrected eDNA rate (β = 0.0011±0.00016, *p* < 10-7), but salmon countsfrom one day prior were only marginally related to any residual variation from the first model (β = 0.00026±0.00015, *p* < 0.09), and salmon counts from two days prior were completely unrelated to residual variation not accounted for by salmon counts from the same day and one day prior (*p* = 0.99).

When pooling 2015 and 2016 data, of the three coho salmon life-history categories (adults, jacks, and nomadic juveniles), adults produced the strongest flow-corrected eDNA signal (β = 0.0059±0.00048, *p* < 10-15), which was 3.5 times higher than that produced by each juvenile fish class (β = 0.0017±0.00058, *p* < 0.004). When accounting for the eDNA signal produced by adults and juveniles, counts of jacks were uncorrelated with flow-corrected eDNA (β = −0.0006±0.0014, *p* = 0.69).

## Discussion

Since the efficacy of eDNA was first demonstrated for the detection of invasive bullfrogs (Ficetola *et al.* 2008), a rapidly growing body of literature has highlighted the efficacy of eDNA for rare species detection (Rees *et al.* 2014; Wilcox *et al.* 2016), has explored the technical aspects of eDNA (Goldberg *et al.* 2016), and has suggested that eDNA holds promise for quantifying the abundance of species (Doi *et al.* 2015; Lacoursière-Roussel *et al.* 2016; Takahara *et al.* 2013; Tillotson *et al.* 2018). The next, and most transformative, technical step for mobilizing the use of eDNA for resource managers is to determine whether, and under what conditions, eDNA can be used to *enumerate* organisms. The possibility of enumerating Pacific salmon as they outmigrate or return to spawn represents a particularly promising application, with large economic and risk-management implications for a multibillion dollar fishery and keystone wildlife resource.

To test the efficacy of eDNA for salmon enumeration, we coupled a complete census of returning and outmigrating anadromous salmon with daily quantitation of environmental DNA. We have demonstrated that flow-corrected eDNA rate:

(1) predicts same-day, daily counts of two species of adult salmon returning into the watershed (Figs. 2, 3) and of one species of outmigrating salmon smolt (Fig. 4),

(2) does not simply accumulate over time, which would have otherwise reflected the total number of salmon that have entered the watershed this season (Figs. 2, 3),

(3) is minimally affected by upstream-rearing juveniles (*Ecology of eDNA*), given that the eDNA from the coho and sockeye fry rearing in Auke Lake appears to settle and/or attenuate prior to reaching lower stream reaches,

(4) is highly accurate at delimiting the phenologies of returning adult and outmigrating juvenile salmon (Figs 2, 3, 4), and

(5) is affected by differential DNA-shedding rates across different life-history strategies and body sizes (*Ecology of eDNA*).

We have also identified several remaining obstacles to straightforward implementation of eDNA for the enumeration of salmon. Most importantly, accurate measures of streamflow are crucial. This is particularly true because pulses of adult salmon immigration sometimes coincide with high streamflow events (Figs. 2-4), and the error in estimating streamflow is exacerbated because the ratings curves that relate river height (the measure that is actually recorded daily) to flow contain more error at extreme values, since extreme-flow estimates are either based on few calibration points or on none at all and just represent extrapolations.

The adult sockeye runs are excellent examples of the importance of obtaining accurate streamflow data (Fig. 2). In 2015, non-flow-corrected sockeye eDNA concentration (‘DNA’ timeline) was highest around 1 July and declined monotonically through the month despite few adult returning sockeye in early July. However, early July was also a period of low stream flow. Only after accounting for stream flow (‘Flow X DNA’ timeline), which included a flood event around 15 July, did eDNA correctly predict the observed sockeye immigration peak on 15 July (‘Counts’ timeline). In 2016, there were three non-flow-corrected eDNA peaks (‘DNA’ timeline), the timings of which very closely matched the three count peaks. However, the first two non-flow-corrected eDNA peaks, in early July, were taller than the third peak, which is the opposite to that seen in the count data (Fig. 2). This occurred because the third eDNA concentration peak, in late July, occurred just as streamflow also rose, diluting the eDNA (‘Flow (cfs)’ timeline). The third eDNA peak’s shape and size more closely matched the count data after flow correction (‘Flow X DNA’ timeline), although the third eDNA peak is still smaller than expected based on the size of the first peak. We hypothesize that the streamflow value that we used to multiply the first day of the third eDNA concentration peak was too low, potentially because it was recorded before most of that day’s flow increase had occurred, causing us to under-correct and thus under-predict. We have informally substituted in the next day’s much higher streamflow value (flow during the third sockeye peak rapidly more than tripled from 6.6 to 23.1 cfs between 23 and 24 July), and the third flow-corrected eDNA peak matches the count data more closely (data not shown).

A second critical consideration for quantifying anadromous fish counts with eDNA is the temporal resolution of an eDNA measurement. As adult salmon move upstream, the signal produced by their shedding of DNA attenuates and is eventually not detectable. Therefore, effective monitoring of anadromous fish with highly variable daily counts requires eDNA to be sampled at least daily. Even with daily sampling, we can imagine that the eDNA signal produced by a medium-sized pulse of fish now could be the same strength as the signal produced by a large pulse of fish that passed by hours ago. This ambiguity sets an upper limit on the accuracy of eDNA for quantifying anadromous fish abundance.

How much the above two *within*-stream sources of error reduce reliability in decision-making depends in part on the level of variation *across* streams. If a single stream, regardless of how accurately it is censused, does not reflect regional escapement sizes, due to variation in salmon abundance across streams, it might be more robust to collect data from many streams (probably only feasible with eDNA), even at a cost of reduced accuracy per stream. Currently, the Alaska salmon fishery does not have enough data to judge this possibility.

A third consideration is that some salmon runs contain a mix of individuals with different life histories. This was particularly the case for coho salmon in 2016, for which jacks were numerically dominant early in the run and a nomadic juvenile coho life history strategy was dominant late in the run. Both nomadic juveniles and jacks were rare in 2015. Jacks and juveniles did not produce levels of DNA concordant with the production by adult salmon (Fig. 3), which introduced error into the relationship between flow-corrected eDNA and coho salmon counts (Figs. 5-6). For unknown reasons, coho jacks produced no detectable eDNA when controlling for adults and nomads.

A fourth consideration is the location of sample collection versus the locations of rearing juvenile salmon and spawning adults. Given our results, salmon enumeration should occur in lower stream reaches, as far as possible from spawning areas that will shed large quantities of eDNA from gametes and decaying fish and from large numbers of rearing salmon fry. It is possible that the presence of the lake upstream of our sampling location facilitated settling or degradation of eDNA, which may have increased the ratio of signal (current salmon moving past the weir) to noise (eDNA from other sources upstream) in our measurements. Similarly, the presence of the weir led to fish released upstream shortly before eDNA sampling.

Implementation in a stream

Finally, noise in enumeration with eDNA can be caused by a lack of primer specificity. Our assays are much more sensitive to sockeye and coho salmon DNA than to non-target salmonids, but there can be non-zero amplification of some non-target DNA. In particular, chinook and coho cross-amplify at low levels (data not shown), which was not an issue in this research because Auke Creek does not have a resident population of chinook salmon (although strays do attempt to enter at the weir). Ensuring good primer specificity to the extant species will help reduce noise in future efforts to enumerate anadromous fish with eDNA.

Pacific salmon are a valuable resource, but their distributed spawning and rearing habitat, due to their anadromous life history, makes monitoring their distribution and abundance a formidable challenge, which consequently injects an unknown but probably non-trivial amount of inefficiency and risk into management. Given the strong observed correlations between daily eDNA samples and fish counts (Figs. 5-6), investment in technology to allow frequent or even near-real-time eDNA quantitation and stream-flow measurement could provide a more accurate and cost-effective means of reducing this inefficiency and risk. This would be especially true if daily eDNA samples from many streams turn out to provide a more accurate estimate of regional escapement sizes than do intensive direct-count measurements at a few streams. However, our results also suggest that using eDNA to estimate fish abundance will require (1) accurate and ideally time-averaged streamflow measures and (2) frequent (at-least-daily) eDNA sampling due to the ephemeral nature of the eDNA signal. On the other hand, this very ephemerality is what makes eDNA such a sensitive correlate of salmon abundance.

Even with a fixed budget constraint, it should be possible for a technician who would otherwise be paid to count fish in a single stream to instead collect water samples from many spawning streams across a watershed. In addition, water sampling could be extended to quantify smolt runs, which are currently only estimated in Southeast Alaska at a small number of index systems. Moreover, because post-sampling filters can be stored in a refrigerator or freezer for many days after sampling, it should be feasible to train and pay a network of citizen scientists to carry out sampling across multiple watersheds. Note also that although our analysis focused on sockeye and coho salmon, the same eDNA sample can be used to detect and/or quantify any number of aquatic species with the development of appropriate assays. Against these potential gains in sampling efficiency and information must be balanced the additional cost of the qPCR assays to be carried out in a dedicated eDNA lab.

Our study is of a single stream in Southeast Alaska. However, it provides strong justification for an expanded effort to sample salmon eDNA over more streams, more species, and more days, both in the streams that currently have weirs, so that a robustly transferable model can be parameterized and validated, and in some of the many streams that are not currently monitored, to test for the possibility that multiple streams sampled daily with eDNA provide more useful information than a few streams counted intensively. The applicability of eDNA to expand monitoring of anadromous salmon to currently unmonitored rivers will depend on the transferability of flow-corrected eDNA rate among streams. It is possible that differences in stream size, morphology, and hyporheic flow will be too idiosyncratic for results calibrated on one weir to be transferable among rivers, thus requiring independent calibration on every river to be monitored. Alternatively, results might be transferable among systems with similar morphology. For example, Auke Creek is a short river course below a lake, which may lead to calibration results that are only transferable to systems with an upstream lake where eDNA settles prior to downstream transport. Given the huge size of the Alaska salmon fishery, even a small improvement in management effectiveness and/or a small decline in the risk of population decline or establishment by alien salmonids could justify the investment in large-scale eDNA calibration tests and an assessment of the efficacy of deploying eDNA to expand the portfolio of streams that can be effectively monitored.

## Acknowledgements

We thank Oregon State University and The National Geographic Society (#9493-14) for funding this work. Thanks to K. Smikrud and S. Pyare for Figure S1. D.W. Yu and C.Y. Yang were supported by the National Natural Science Foundation of China (41661144002, 31670536, 31400470, 31500305), the Key Research Program of Frontier Sciences, CAS (QYZDY-SSW-SMC024), the Bureau of International Cooperation project (GJHZ1754), the Strategic Priority Research Program of the Chinese Academy of Sciences (XDA20050202, XDB31000000), the Ministry of Science and Technology of China (2012FY110800), the University of East Anglia, and the State Key Laboratory of Genetic Resources and Evolution (GREKF16-09) at the Kunming Institute of Zoology. D.A. Tallmon was supported by North Pacific Research Board project #1710. The findings and conclusions in the paper are those of the authors and do not necessarily represent the views of the National Marine Fisheries Service. Any use of trade, firm, or product names is for descriptive purposes only and does not imply endorsement by the U.S. Government.

## Data Accessibility

The R scripts and data for analyses are at github.com/dougwyu/2014_2015_2016_Auke_qPCR, and on Dryad doi: 10.5061/dryad.94d37g3

## Author Contributions

TL, DWY, DAT, and CYY conceived the research and designed the experiments. DB, JJ, SCV, and JRR collected field samples. JA and CYY performed laboratory analyses. TL and DY performed the statistical analysis. TL, DWY, and DAT wrote the manuscript, with comments from all other authors.

## Supplemental Information for

**S1.**
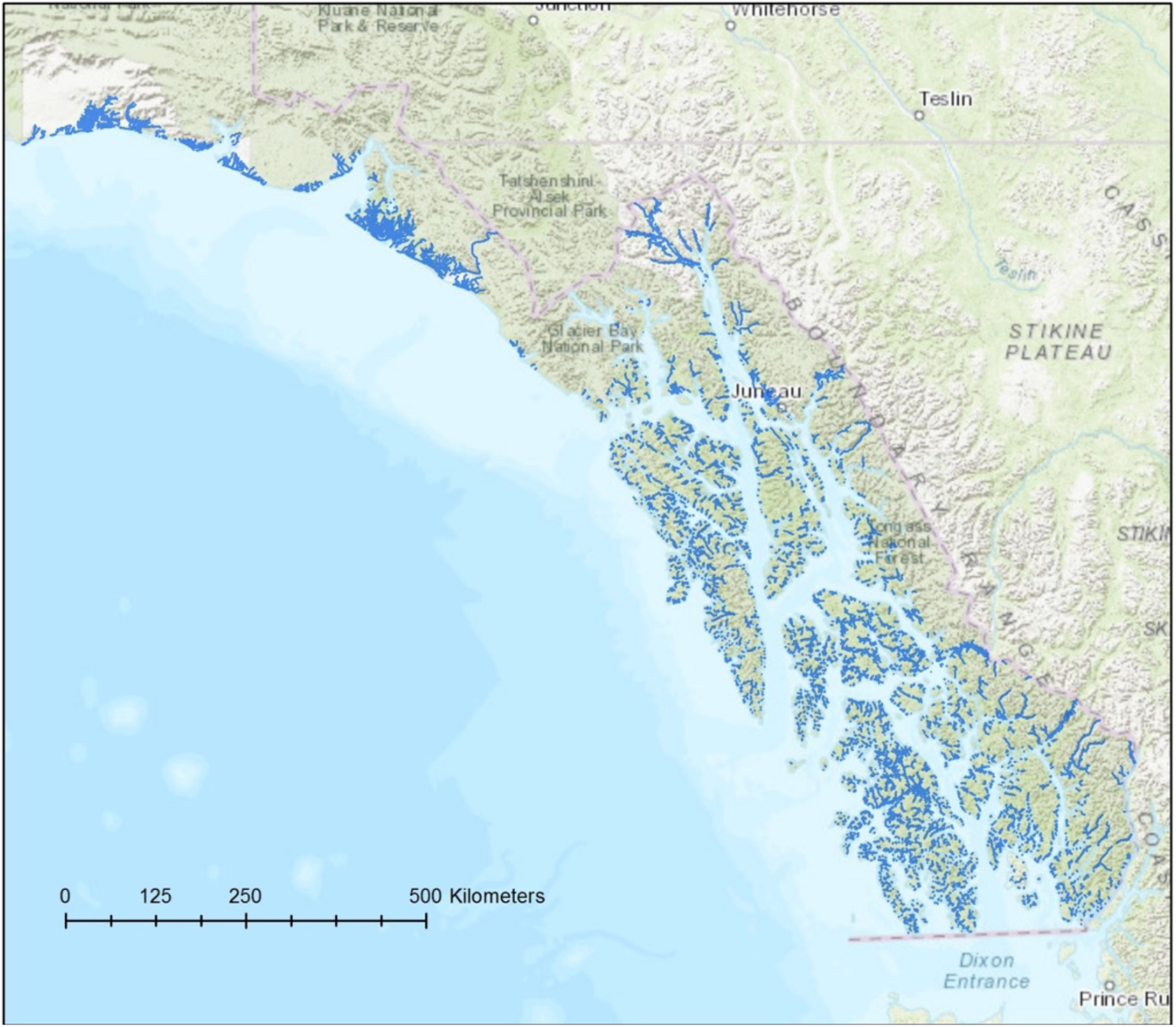
Salmon spawning streams in Southeast Alaska indicated by blue shading.

**S2.**
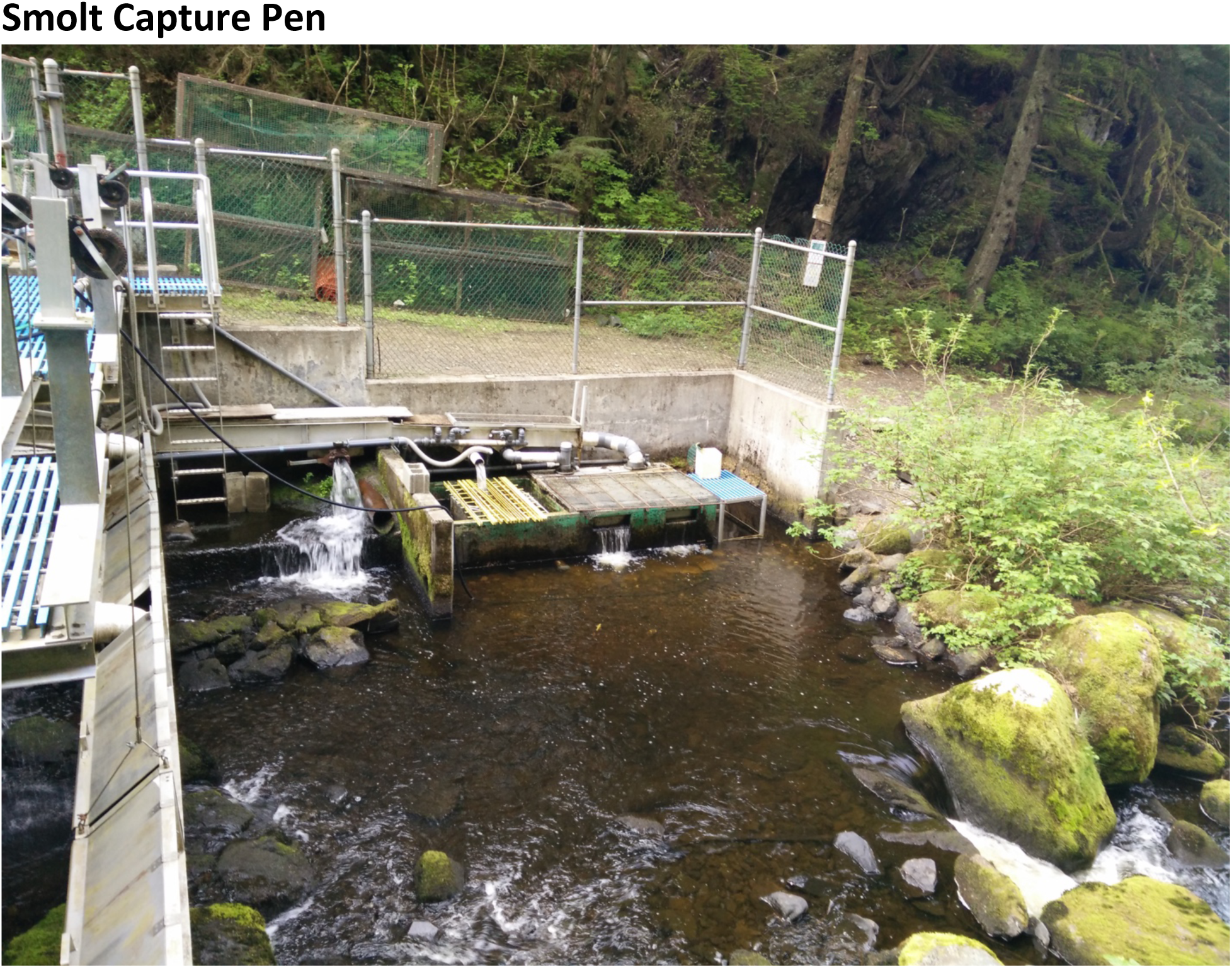
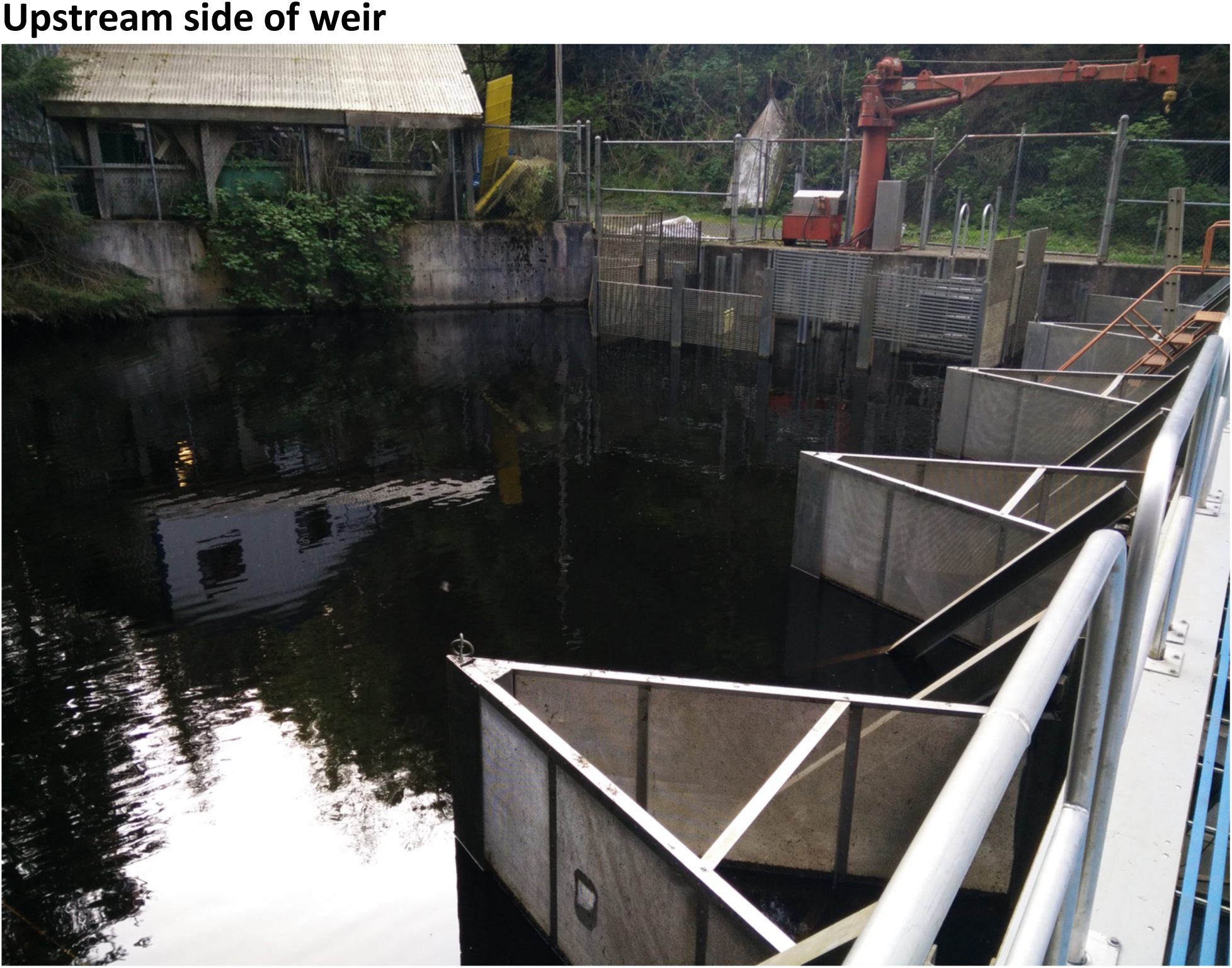
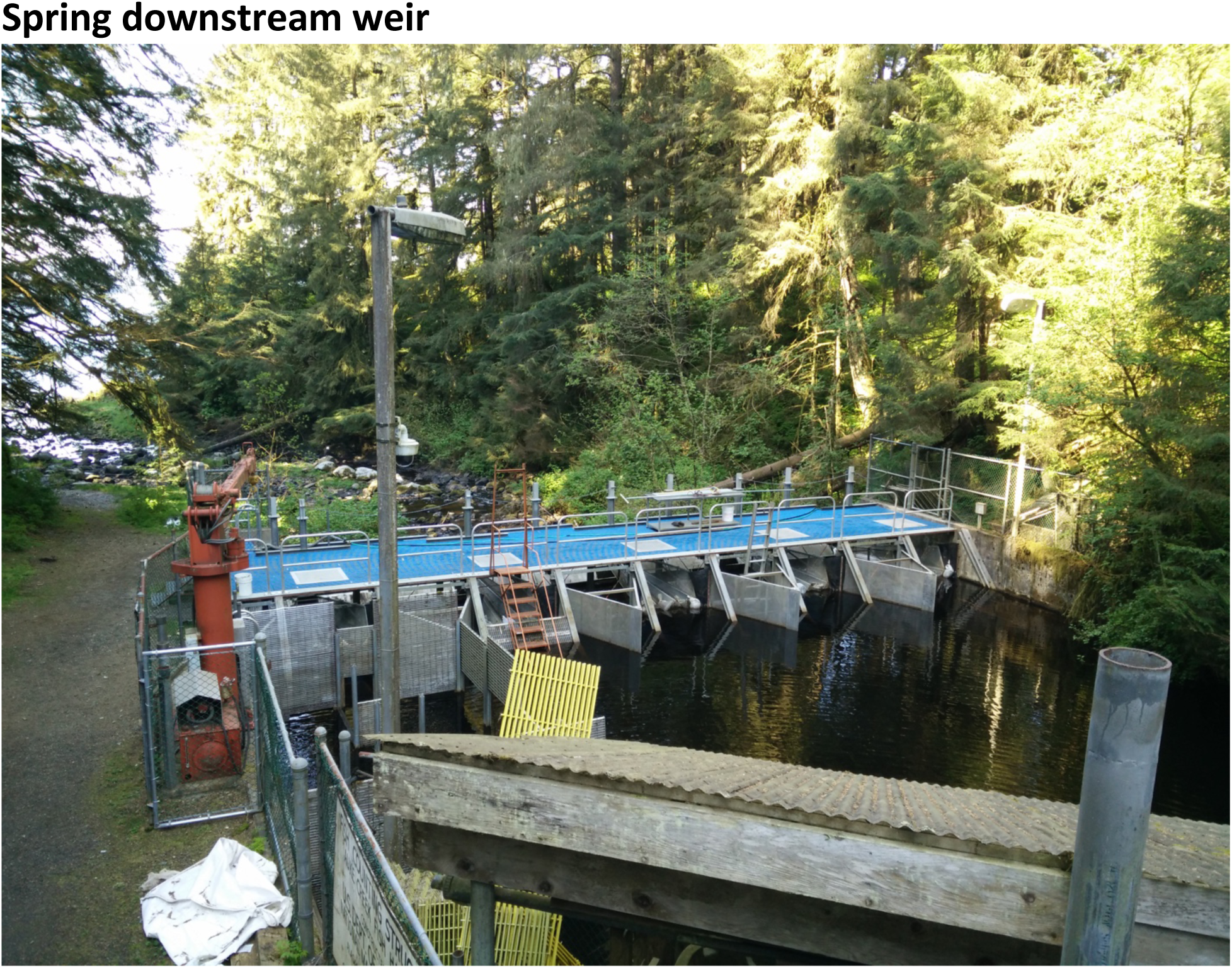
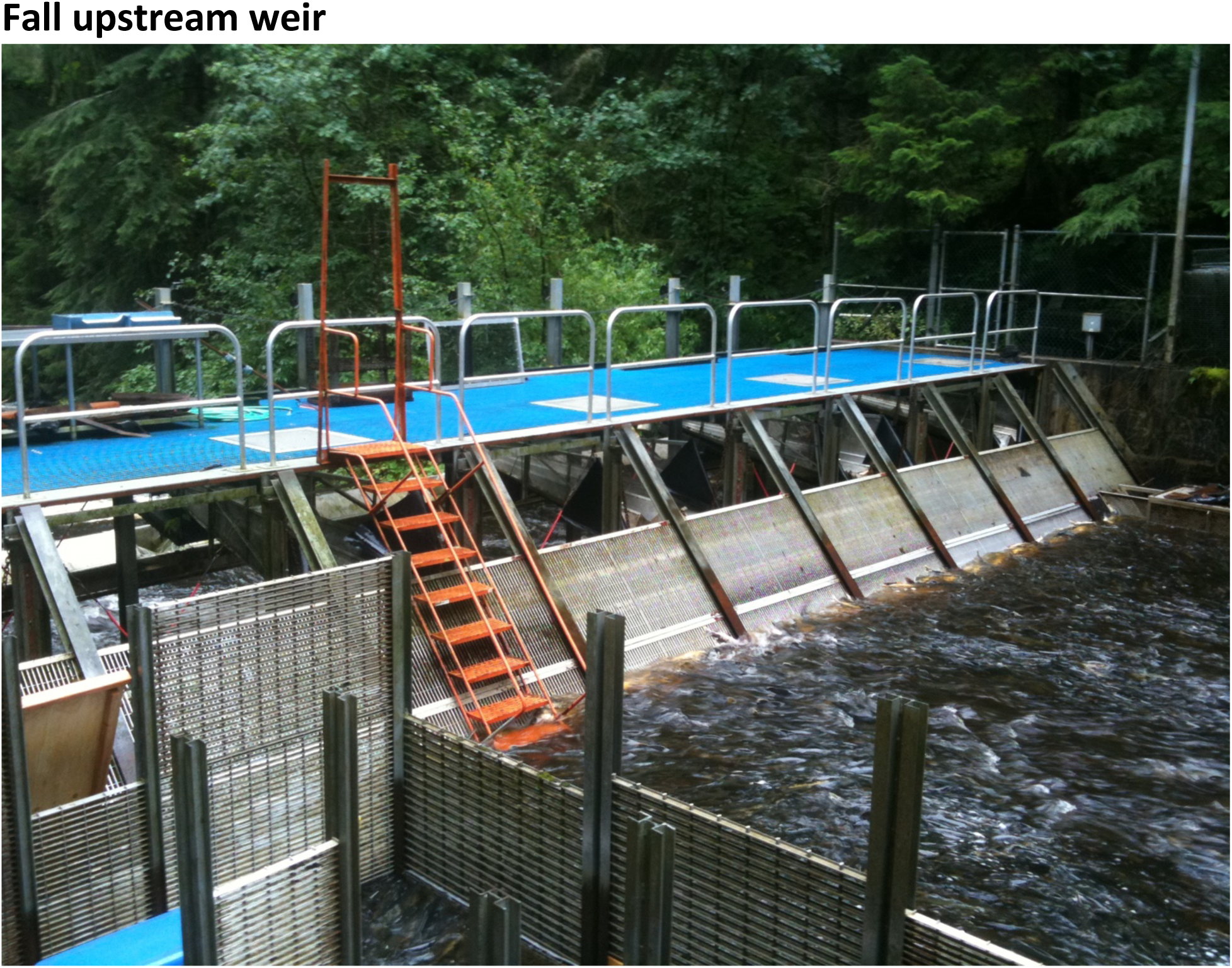
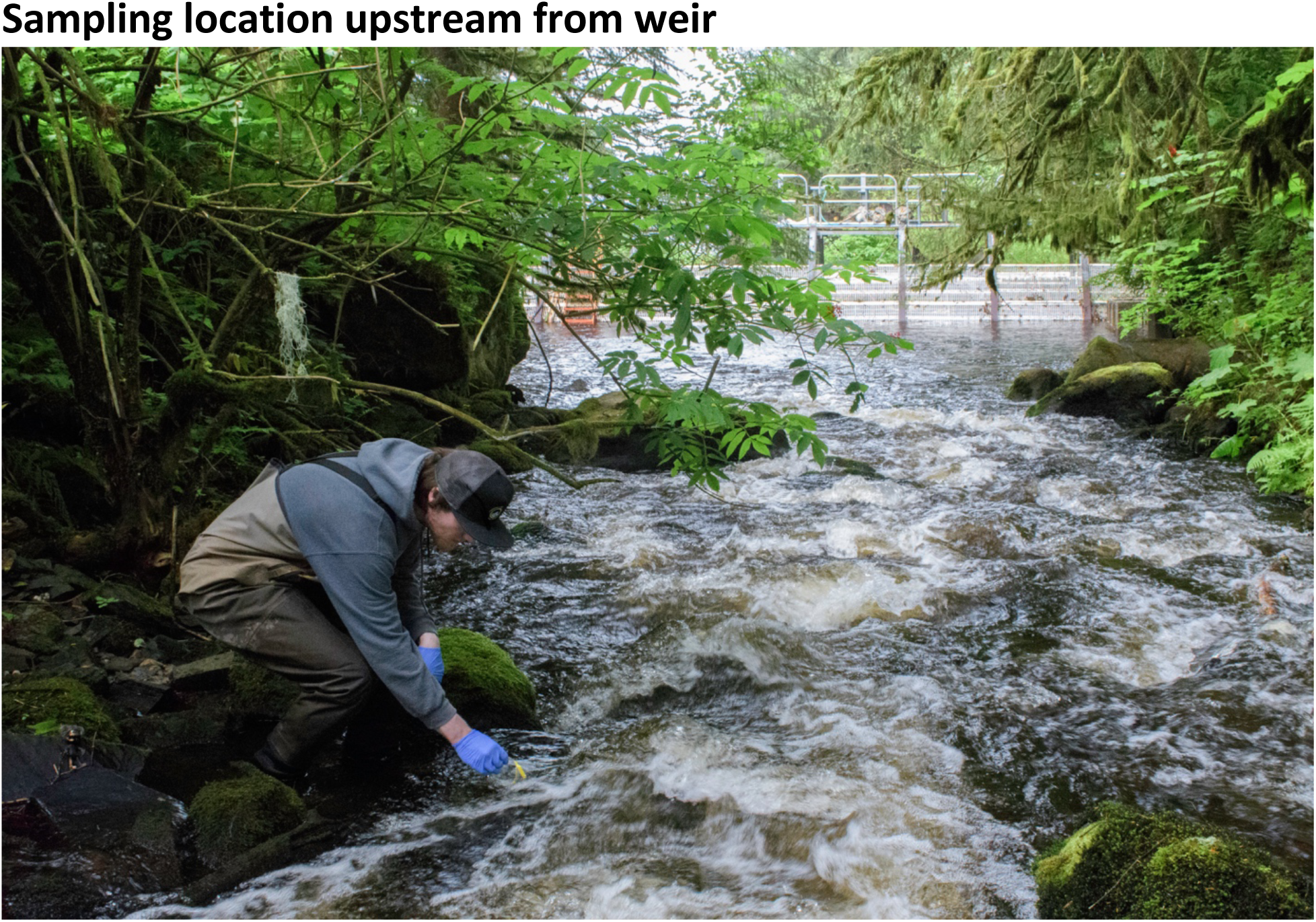
Photos of weir structure

**S3. Results including outlier day with pulse of jacks**

**Figure S3.1.**
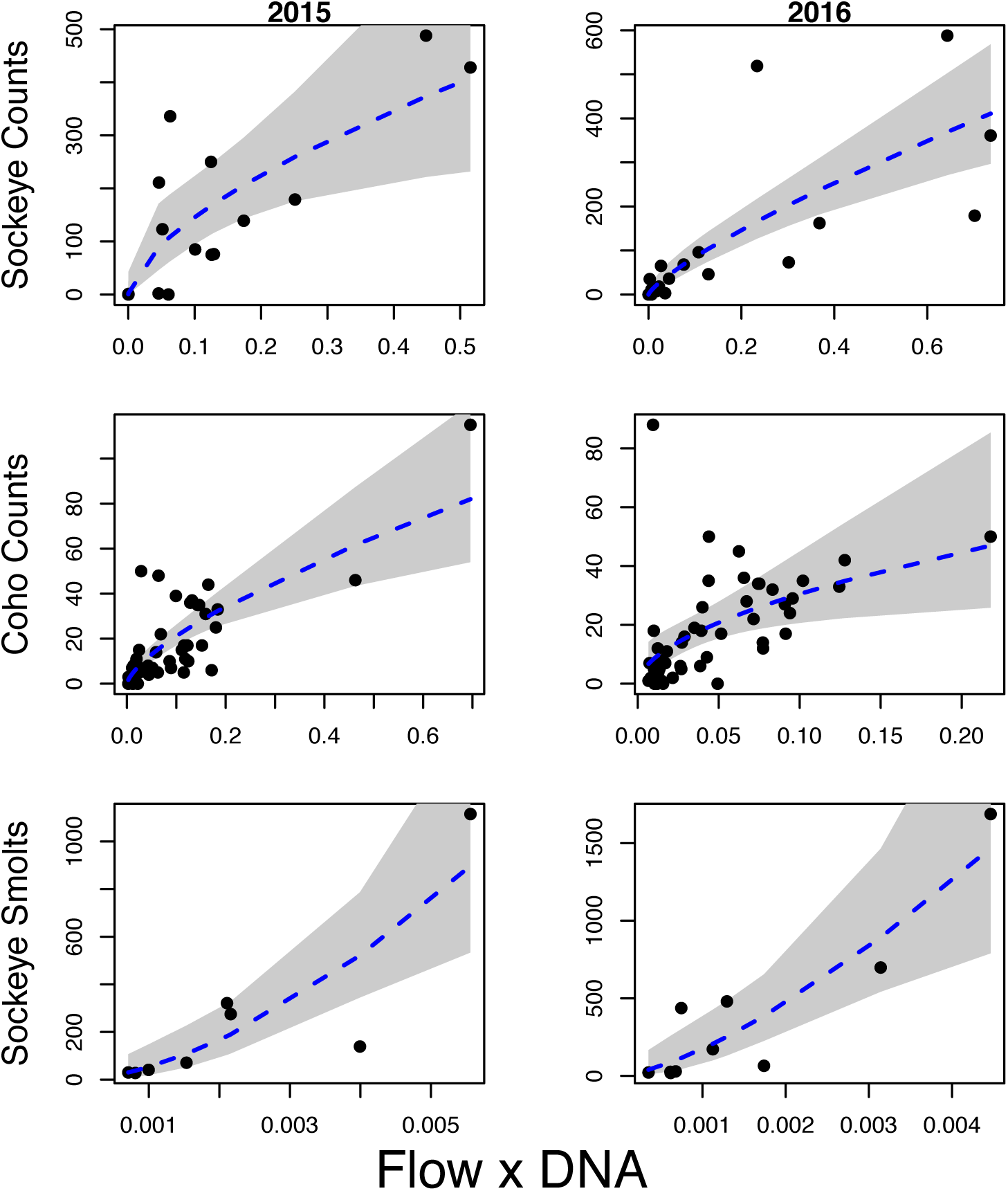
Results of quasi-Poisson regression models relating the natural logarithm of flow-corrected eDNA concentration to adult sockeye salmon counts (2015: p=0.008, 2016: p<2e-8), total coho salmon counts (2015: p<2e-8, 2016: p=0.002), and counts of sockeye salmon smolts (2015: p=0.004, 2016: p=0.003). Gray shading denotes the 95% confidence interval.

**Figure S3.2.**
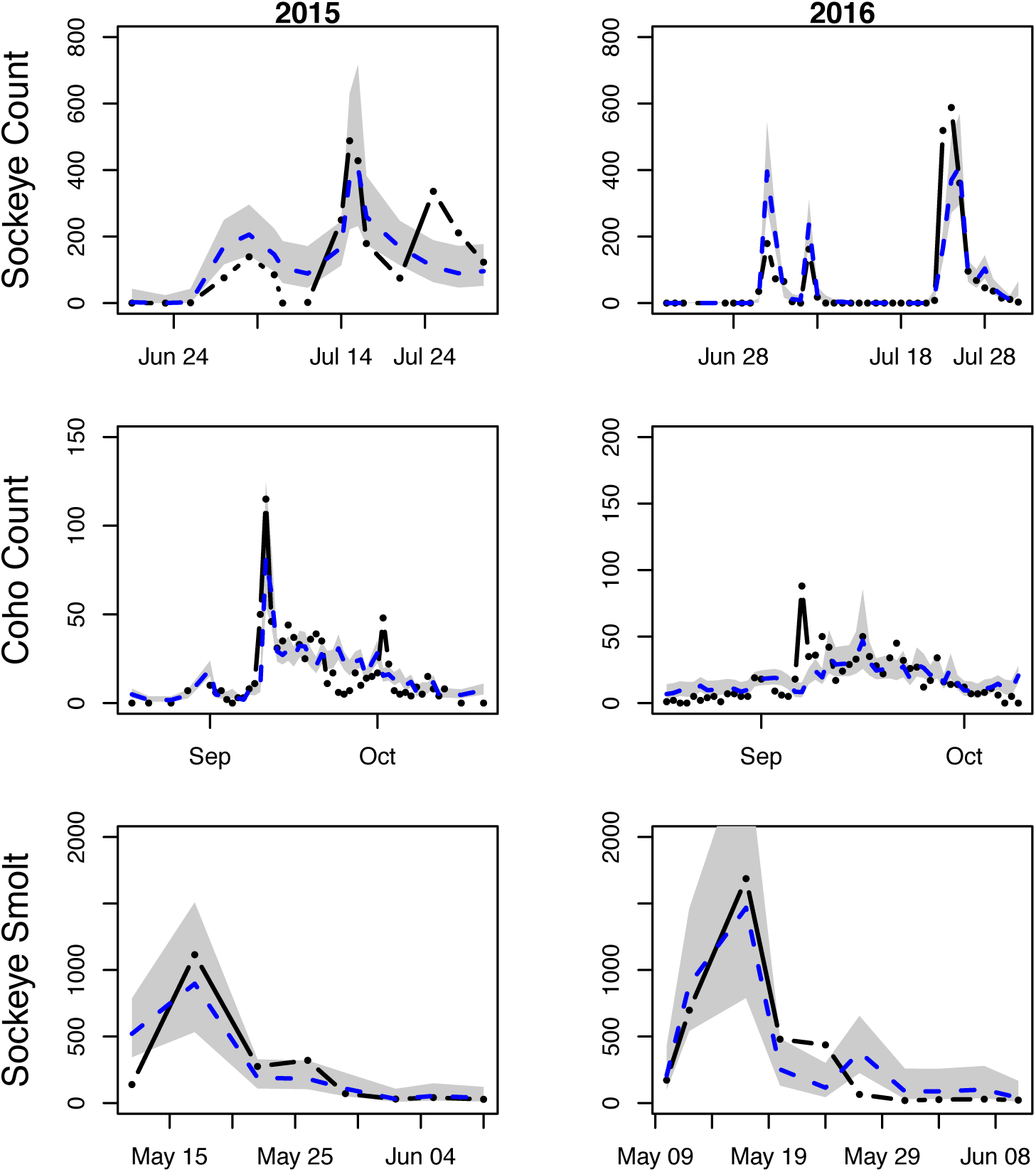
Counts of adult sockeye salmon, total coho salmon (including all life history strategies and the single outlier in 2016), and sockeye salmon smolts (black dots) and the predicted number of counts based on the flow-corrected eDNA concentration predictor in the quasi-Poisson regression model (blue dashed lines). Gray shading denotes the 95% confidence interval.

